# A Multiscale Model of Collective Decision-Making in Hybrid Aspen Tree Tissues Describes Bud-Dormancy Break

**DOI:** 10.64898/2026.02.04.703817

**Authors:** Hannah Dromiack, Swanand Khanapurkar, Rachel Phillips, Tatiana de Souza Moraes, Gwendolyn V. Davis, Shashank Pandey, Bibek Aryal, Aswin Nair, George W. Bassel, Emmanuelle M. Bayer, Rishikesh P. Bhalerao, Sara I. Walker

## Abstract

The mechanisms underlying cellular coordination within tissues remain enigmatic. Most models focus on interactions between just two levels of organization - cell and tissue - and do not leverage data across deeper hierarchies that best represent living processes, with many spatial and temporal scales interacting. Integrating many scales, from molecular to cellular to tissular to organismal to populational, may be necessary to fully elucidate tissue function, especially in cases of sparse data at each level. Here, we investigate multiscale, robust regulation of tissue-level decision-making, using experimental studies of cold induced dormancy release in terminal buds of hybrid aspen trees as our case study. We develop a network model of terminal bud meristematic tissue, incorporating expression data from a key cold induced regulator gene, FLOWERING LOCUS T (FT1), which controls bud dormancy release, combined with data on variability in cell-to-cell communication controlled by FT1 mediated regulation of plasmodesmata. The model can explain dormancy breaking under constant temperature, but not variable temperature. We introduce constraints from organismal-level data and show how the presence of coordinated cellular interactions within individual plant tissues is necessary to reproduce data of population-level statistics. Our findings demonstrate how mechanisms of tissue function may be better constrained when data are used across more scales. They also hint at potential tantalizing new insights such as how tissue function might not be solely dictated bottom-up from molecular interactions, but also top-down from constraints imposed by the organismal and population context. Both implications illustrate the critical importance of incorporating cross-scale information processing in modeling biological decision-making.

**Significance Statement:** Biological hierarchies involve decision-making mediated via nested feedback loops. Data-informed modeling of this hierarchal complexity remains challenging. Here, we leverage unique features of plant biology - stationary growth, prolonged decision-making, and physical structure - to study multiscale dynamics determining cellular mechanisms of bud dormancy breaking in aspen trees. We examine how tissue function can be driven bottom-up from gene regulatory networks and top-down from organismal population-level statistics. Modeling experimental data collected at genetic, cellular, and organismal levels, reveals how population-level data allow constraining mechanisms of cellular coordination within individual plants when cellular data are sparse. Our findings demonstrate how multiscale methods can combat data sparsity and suggest new ways to study cellular coordination within organisms could be dictated by organismal population-level constraints.

## Main

Collective decisions arise when group-level outcomes cannot be attributed to any single individual. In biological systems, collective decision-making spans multiple organizational levels—from how cells coordinate during morphogenesis [18, 55, 56] to how social insects select new nest sites [11, 34, 54]. Most studies on collective decision making generally address only two levels—individuals and their collective, although it is evident that collective behaviors occur across hierarchies of organization [17, 30, 47, 52, 59]. For example, cells form tissues, tissues form organisms, and organisms form populations, each exerting both upward (emergent) and downward (constraining) causal influences [13, 36, 50].

The concept of hierarchical organization has deep roots in the study of biology; one cannot detach any one level (say the cells) from another (the multicellular individual) and still accurately capture the full picture of the detached level because living systems are so deeply integrated across these hierarchical scales [15, 41]. Yet, most models reduce systems to two levels, overlooking how lower- and higher-order dynamics interact. Successful examples include models group motion of swarms or of fish schools [8, 9, 53]; these models account for how individual-organismal behaviors result in emergent group-level patterns. However, expansion beyond this paradigm has shown that social behavior often reflects group-level constraints on individuals [27, 37, 38], and more recently bird flocking depends on neuroanatomical mechanisms within individual birds as well as group-level dynamics [49]. While the existing body of collective behavior is rich [2, 7, 16, 21, 22], expanding collective behavior models to include multiple nested levels represents a frontier with potential to reveal mechanisms that cannot be uncovered in shallower hierarchies (e.g. two scales).

All living systems are nested multi-level systems. However, plants represent one of the most tractable model systems for studying the deep hierarchies of biological systems due to their stationary disposition, prolonged decision-making timescales, and mappable tissue structure. Plants have no centralized decision-making system, necessitating a need to study collective coordination across scales to understand their behavior [10]. Plants make up nearly 80% of the biomass on Earth [5]. They cycle and sequester carbon, provide heat sinks, are responsible for global cooling, and are keystone species for forest and grassland ecosystems across Earth [48]. In this study, we focus on hybrid aspen trees of genus *Populus*, a model for studying perennial trees [25, 48, 58].

Perennial trees such as hybrid aspen undergo cycles of growth and dormancy, crucial for adaptation to seasonal changes [33]. Prior to winter, growth in shoot apex is suppressed and the shoot apical meristem (SAM) and growth arrest leaf primordia are enclosed within a protective apical bud structure. Subsequently after growth arrest dormancy is established in the apical bud and for growth to resume in the spring, dormancy is broken by prolonged exposure (2 – 6weeks) to low temperature (4 – 8°C). Transitioning from dormancy to active growth is an expensive process for a plant, and an irreversible decision [6]. Therefore, plants must be able to integrate environmental information to determine the optimal time to risk growth and, due to the magnitude of the commitment, be robust to error [12]. Thus, understanding the mechanisms for how dormancy is broken, given specific environmental conditions (e.g., temperature, humidity, salinity, etc.), is critical to many areas of study from fundamental biology and ecology, to mitigating the economic impacts of climate change.

Dormancy release (or breaking) primarily manifests at the cellular level via FLOWERING LOCUS T (FT1) [57], **Figure 1A**, and its regulation of intercellular channels known as plasmodesmata (PD), **Figure 1B**, yet it is also known to exhibit tissue-to-population effects, **Figure C-E**, in the form of population-level bet-hedging [3, 20]. Bet hedging occurs when populations have heterogenous responses, increasing the probability of some individuals surviving over others in the face of uncertain environmental variability [1, 20]. During dormancy establishment, cell-to-cell communication via PD is blocked by deposition of callose [31]. Conversely, low temperature triggers dormancy release by unblocking PD by downregulation of callose. Recent studies have identified a key role for FT1 gene in cold induced dormancy release [42]. In hybrid aspen, FT1 expression is induced by cold and scales with exposure to cold. FT1 mutant variants fail to undergo dormancy release. Importantly, FT1 facilitates dormancy release by opening of PD to restore cell-cell communication within the bud tissue and interestingly, FT1 itself moves via PDs [31]; thus, presumably FT1 promotes its own movement generating a feedforward loop. Moreover, recently it was shown that bud dormancy also displays population-level bet-hedging strategies as revealed by the heterogenous response of bud dormancy release in genetically identical plants, which is enhanced under variable conditions [1, 3, 20]. Consequently, bud dormancy release offers a highly useful experimental system to investigate multilevel control of collective decision making.

**Figure 1.**
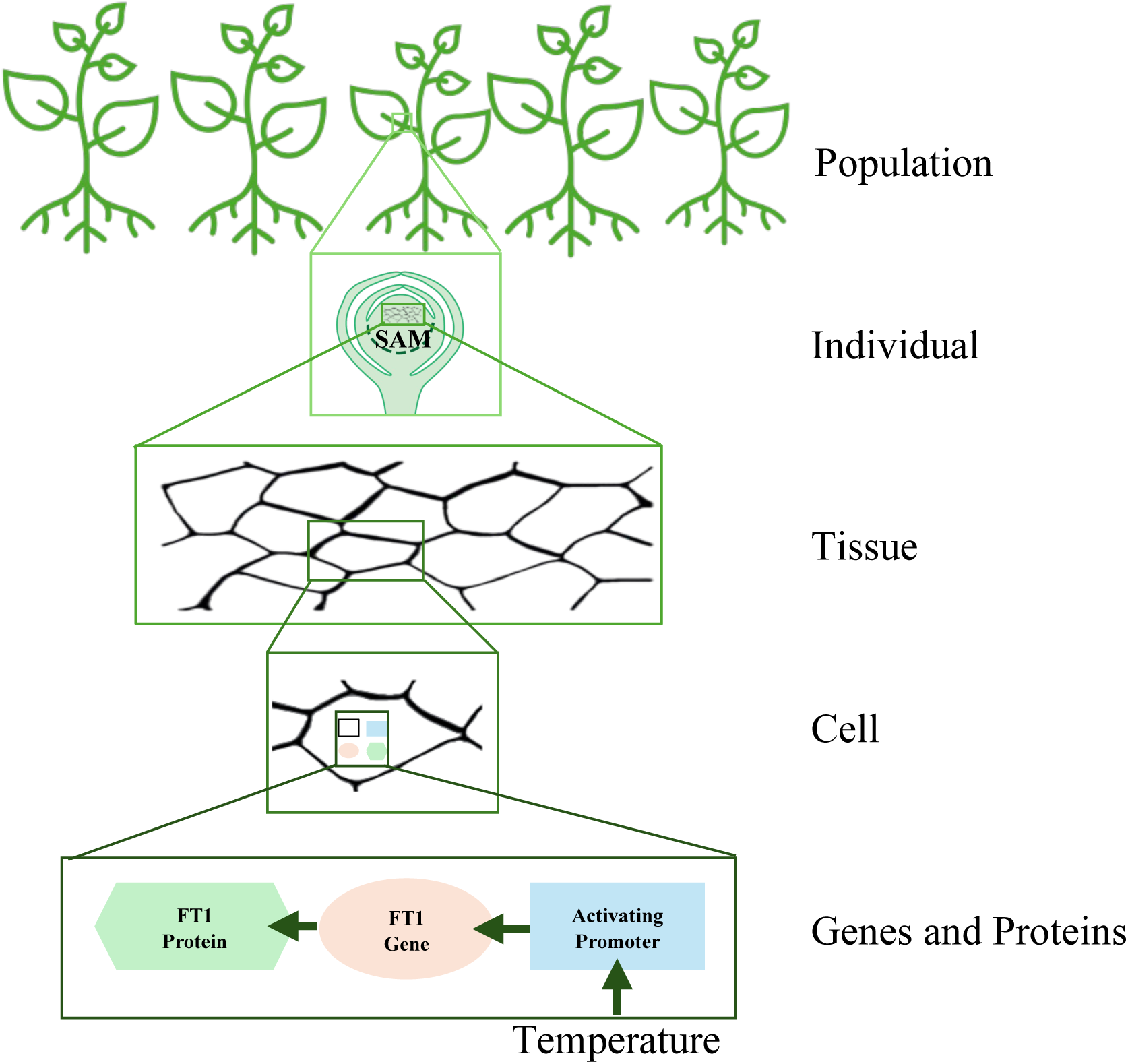
Levels of organization from subcellular genetic circuits to populations of Hybrid Aspen Trees, which could influence tissue-level decision making. The lowest level is a gene regulatory circuit responsible for FT1 expression in response to environmental temperatures. Each cells contains the gene regulatory circuit and the presence of FT1 drives dormancy breakage by promoting cell-to-cell communication by modifying PD dynamics. Tissue structure and PD states (open or closed) dictate the movement of proteins across the tissue. Cellular interaction networks are defined from the cellular topology of the Shoot Apical Meristem (SAM) tissue and inform structures of the simulated tissues. For each individual plant, above ground growth occurs in the terminal bud which shows macroscale changes visible to an observer when dormancy is broken and growth begins. Finally, populations experiencing the same conditions display variation in dormancy release despite being genetically identical.

In what follows, we leverage the genetic and cell biological control of bud dormancy regulation, combining experimental data from gene expression, PD response and dormancy release data to model how cellular decision-making dynamics drive bud dormancy breaking in *Populus spp.* tissue. In systems where data is sparse, multiscale analyses can connect data types across different scales via mechanistic models that are mutually co-constraining. This allows identifying hypotheses for collective decision-making mechanisms that can then be iteratively tested. We demonstrate collective cellular coordination may be necessitated by population-level outcomes, illustrating how integrating bottom-up and top-down processes to constrain tissue function can reveal collective decision-making mechanisms that cannot be easily deciphered by single-scale analyses.

## Results

### Data-Driven Multi-scale Modeling of Tissue-level Decision Making in Hybrid Aspen Trees

We use experimental data including cellular tissue structure (including PD state distributions at cell interfaces), gene expression rates for FT1 protein in cellular tissue, dormancy states of individuals under study, and time of bud break for all individuals within a population [42]. PD states for wildtype tissues comes from visual inspection of raw data obtained by confocal microscopy using callose immunofluorescence. The meristematic region tissue map is converted to a corresponding network representation. Here, nodes correspond to cells, and edges to cell walls, which contain some number of PDs, see **Supplement 2**. States of the nodes and edges are treated as Boolean variables: cells (nodes) can either be expressing FT1 or not (1 or 0) and PDs in cell wall (edges) can be opened or closed (1 or 0). Cell walls in a real tissue can have a range of PDs on each wall, with any percentage open; for the Boolean variable representation, an edge is considered open if any single PD on that wall is open allowing adjacent cells to communicate.

### Plasmodesmata Permeability Clustering and Model Cell Wall Dynamics

Meristematic tissue is a lattice-like network which is structurally static throughout the decision-making process of breaking dormancy: cells do not elongate, divide, or alter neighbors [46]. We hypothesize that how cells are arranged relative to one another in the meristematic tissue structure aids in the decision to break dormancy by limiting the speed by which FT1 expression spreads by the presence of physical barriers (i.e., the cell walls containing PDs) and the size of the tissue that FT1 must permeate through. Modelling the tissue as a network, we tested this hypothesis by comparing the wildtype tissues structure to connected Erdős Rényi (ER) random graphs conserving the distribution of nodes and edges [14]. If the wildtype tissue structure were indistinguishable from random, it would suggest that physical barriers and tissue scale organization have little influence in dormancy release, and subsequently that tissue structure is not selected to aid in this process.

Using a standard suite of network statistics [24], we found that variation between wildtype and random networks is evident in average shortest path, network diameter, and clustering coefficient, indicating that the tissue structure has cellular organization with non-random attributes [40], **Sup Figure 4**, (for full suite see **Supplement 5)**. The presence of cellular organization could inform communication processes that may play a role regulating how FT1 travels throughout the tissue. To further investigate this, PDs states from wildtype tissue samples are overlaid onto the networks, as either open or closed edges, to determine how the tissue structure may dictate FT1 transport and therefore play a role in dormancy breaking. **Figure 2A** overlays the data visualization of PD activity on the original cross-section tissue sample image (after 4 weeks of continuous cold) [42]. **Figure 2B** shows the network representation including corresponding edge states. Sample sets represent the dormant, 1-week, 2-week, 3-week, and 4-week marks in the continuous cold treatment. As dormancy is broken, the number of cell clusters connected via OPEN edges tends toward 1, **Figure 2C**; this is consistent with how when dormancy breakage is decided, it will become a whole tissue response. Finally, cellular clustering increases throughout the decision, **Figure 2D**, suggesting that adjacent cell wall PDs open in clusters. Clustering within the spatial distribution of PD activity suggests local coordination dynamic among cells with respect to a cell state change to “ON” for FT1.

**Figure 2.**
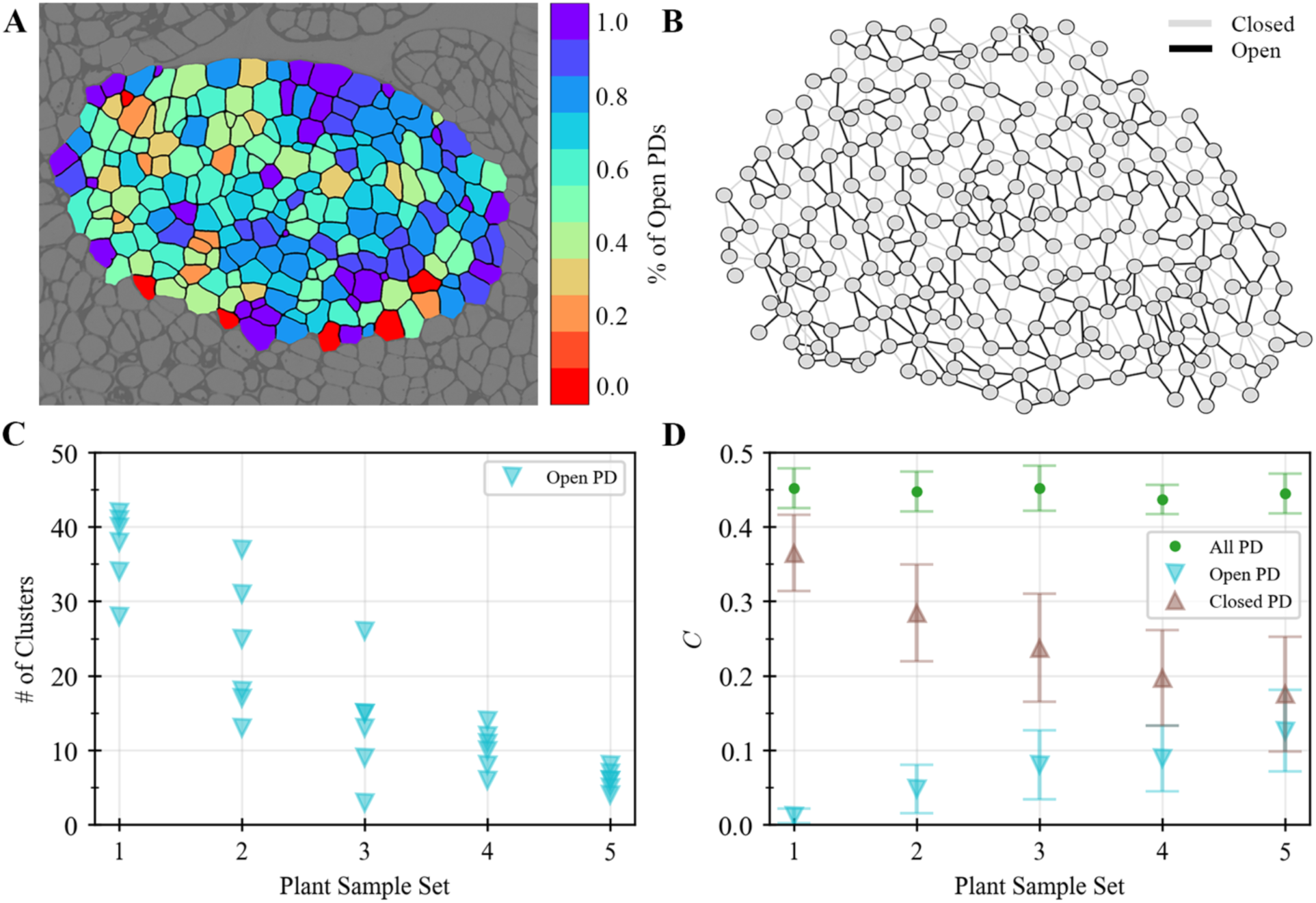
Clustering analysis of wildtype plasmodesmata activity across the tissue. **A)** Cross-section of a tissue sample colored by the percentage of opened plasmodesmata (PD) for a given cell at 672 hours of continuous cold treatment. **B)** A network representation of the cross-section tissue sample shown in A), where nodes represent cells and edges represent cell walls. **C)** Number of clusters connected via open edges throughout the continuous cold treatment, where the network increases in connectivity toward a single large cluster as dormancy is broken. **D)** Clustering coefficient throughout breaking bud dormancy for three representations of the PD networks. Wildtype is the whole tissue structure regardless of PD state in cell wall. Open considers only the portions of the wildtype network with open edges and closed is the converse. The clustering coefficient of cell walls connecting cells does not change across the samples and increases for open walls.

Known causal connections between the expression of FT1 and the opening of PD have been identified previously [42]. Alongside a non-random structure, together this suggests the following dynamic for simulated PD opening, i.e., network edge activation in the model. For an edge to activate in cold temperatures one or both connecting cells must be expressing FT1; if only one is expressing, the edge has some probability 𝑝_𝑒_ of opening, see Methods and/or **Supplement 2**. The parameter 𝑝_𝑒_ is derived directly from experimental data utilizing a least squares algorithm for an exponential fit on the global amount of PD activity in continuous cold treatment (𝑝_𝑒_ = 0.0012, 𝑅^2^ = 0.89), see **Supplement 4: Sup Figure 3**. Finally, the observed cellular organization, cellular connection clustering, and FT1 movement through PDs can inform cellular state dynamics, specifically a cell’s decision to activate FT1 expression. Decentralized systems such as bacteria [39] and social insects [45] utilize quorum sensing to coordinate their behaviors and act in unison, furthermore it is a common mechanism for collective decision making across species and system architecture [51]. Therefore, we present a simple quorum majority rule for cell gene expression state (e.g., if a certain ratio of neighboring cells, 𝑛_𝑐_, are expressing FT1, change state), to test how local coordination might be playing a role in tissue-level decisions.

### An Integrated Two-Level Model of Gene Expression and Tissue-Level Dynamics

The induction of FT1 expression by cold is crucial for dormancy release [3, 42]. To determine potential mechanisms for the induction of FT1 expression by cold temperatures, we test three hypotheses: (1) stochastic activation, (2) a local quorum rule, indicating nearest neighbor communication and quorum sensing within the tissue, or (3) a combination of the two. While there are potentially many mathematical models that could be explored to explain this process, stochastic activation and the quorum rule were chosen to map directly to known mechanisms within the tissue. For example, FT1 is responsive to temperature and therefore the simplest model for this would be some form of thermal stochastic activation. Furthermore, stochastic activation is a common mechanism in gene activation [35, 44] and provides the means to transition cells into an active state at the start without the need for cellular interaction. Also, interestingly, cold regulation of FLC, (a repressor of flowering) in vernalization (a process that shares certain similarities with dormancy release) also involves stochastic control [4]. The alternative mechanism of quorum rule was selected because FT1 is a direct regulator of PD (thus PD opening serves as an output of FT1 activation) and experimental data shows that PD opening is increasingly clustered in response to cold, correlating with increased FT1 expression. Moreover, FT1 is mobile and presumably moves via PD between cells and therefore once PD open, FT1 can diffuse between neighboring cells creating nearest neighbor interactions typically observed in quorum sensing, a common mechanism found across diverse biological systems.

First, we define stochastic activation given cold temperatures as:

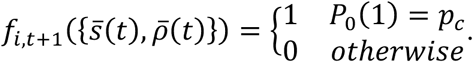

That is, the state at 𝑡 + 1 for the 𝑖^𝑡ℎ^ cell has some probability 𝑝_𝑐_ of activating, regardless of the FT1 expression states of its direct neighbors, 𝑠̅(𝑡), and associated state of their cell wall PD openness, 𝜌̅(𝑡), at some time, 𝑡.

Quorum sensing via local cell-to-cell interaction governed by a quorum rule is modeled as:

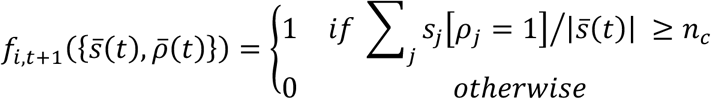

Here, the state at 𝑡 + 1 for any 𝑖^𝑡ℎ^ cell is determined by the number of the nearest neighbor cells which have FT1 expression states equal to 𝑠_j_ = 1, and by the associated state of their cell wall PDs being open, 𝜌_j_= 1, normalized by the total number of neighboring cells at some time, 𝑡. If this value is above a threshold value, 𝑛_𝑐_ (fit to experimental data), then the cell activates. 𝑛_𝑐_ sets the threshold for how many nearest neighbors must be expressing and communicating (i.e., have an open cell wall) for a cell to begin expressing. 𝑝_𝑐_ sets the probability that a cell will stochastically begin expressing when cold temperatures are present.

Simulated FT1 expression and simulated ratio of cell walls with open PD are displayed in **Figure 3A** and **B**, respectively. **Figure 3C**, shows a sweep across the parameter space, with only the final state for the simulated cells and cell wall PD are plotted. These data reveal that both (stochastic activation and local coordination via quorum rule) mechanisms do display some capacity to partially recreate experimental data, either with respect to gene expression rates or PD openness. While the dynamics of cell wall PD openness is not directly determined by either 𝑛_𝑐_ or 𝑝_𝑐_, they are indirectly affected by the number of “ON” cells, which in turn are activated based on cell wall PD dynamics. **Figure 3 top row** displays the results for local coordination via quorum rule, both over the entire transition period and at the final state (four weeks) across different 𝑛_𝑐_ values. Here, it is evident that no single value of 𝑛_𝑐_ can reasonably recreate the FT1 expression data, nor the ratio of cell walls with open PD seen in the wildtype data. Looking solely at the final state (672hrs) there is an evident phase transition at 𝑛_𝑐_ > 0.2, this occurs due to insufficient number of initially ON cells (i.e. expressing FT1), suggesting that above this the quorum rule does not have a prominent role. Below 0.2, at 𝑛_𝑐_ ≈ 0.15 the model with quorum rule can reproduce the final state of the experimental data, but then it undervalues the number of cell walls with open PDs across the duration of the decision. **Figure 3 bottom row** displays the stochastic activation case; again with possible alignment to experimental data, though node states (FT1 expressing cells) and edge states (cell walls with open PD) cannot be simultaneously correlated to experimental data with any one 𝑝_𝑐_value. Thus, neither the stochastic nor the local quorum rule mechanism on its own is sufficient to recreate all the experimentally observed dynamics of FT1 expression or PD openness at tissue level.

**Figure 3.**
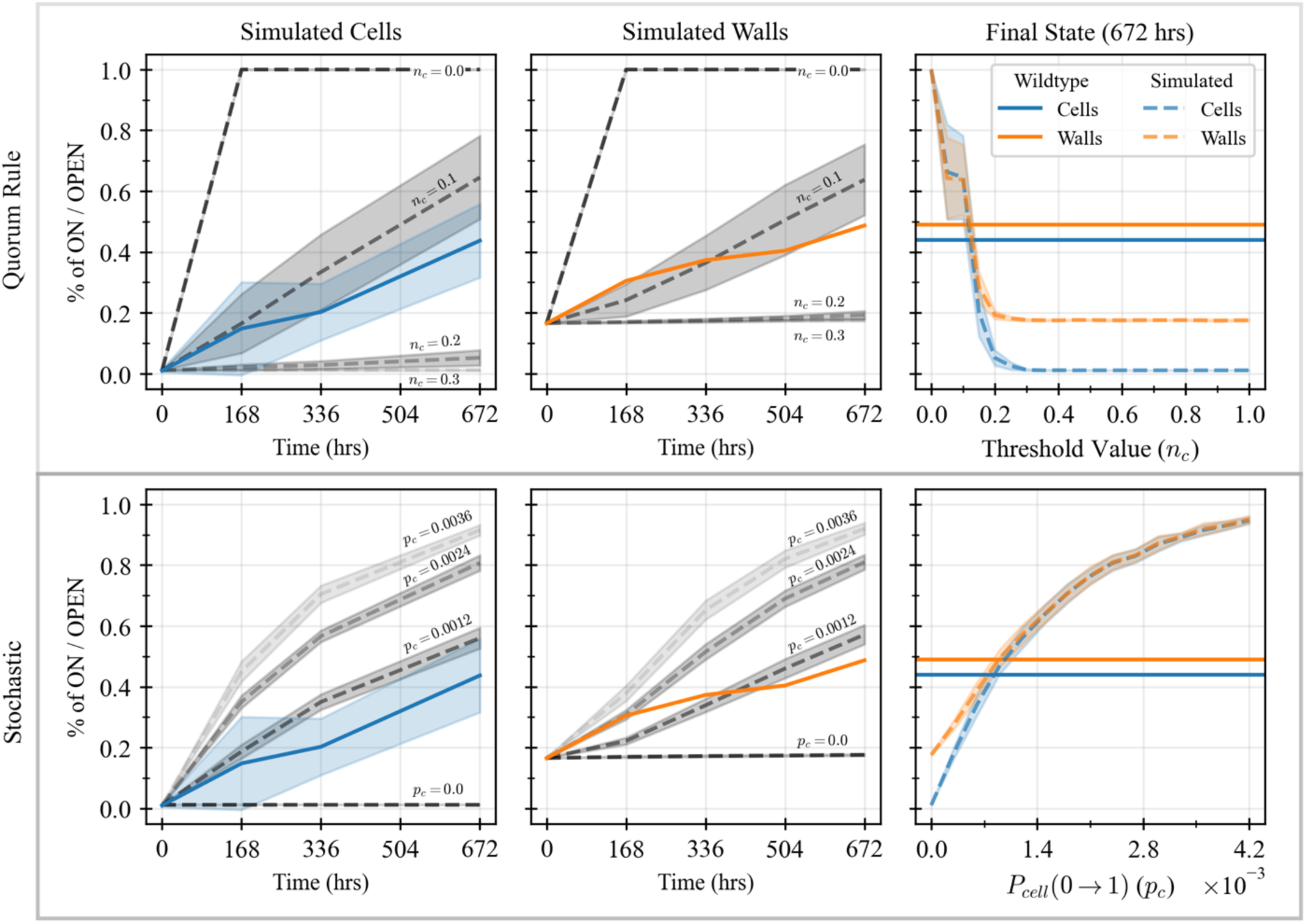
Cell and wall states for simulated tissues using stochastic or collective decision-making rules in continuous cold treatment. **Top row:** Quorum rule for cellular activation. **Bottom row:** Stochastic activation of cellular gene expression. **Left column:** Simulated node FT1 expression levels and experimental FT1 gene expression levels are presented in blue. Middle column: Simulated cell wall with open PD (i.e. communication between these cells is possible), experimental cell wall with open PDs is presented in orange. **Right column:** final expression levels for nodes (cells expressing FT1) and edges (cell walls with open PD) (dashed blue/orange respectively) at four weeks across a range of values for each parameter. While the stochastic process can reproduce the cell activity, it cannot generate the edge dynamics. Given only a stochastic process, the tissue expression levels asymptotically approach one as 𝑝_𝑐_ grows. With only a quorum rule present there is a clear phase transition at 𝑛_𝑐_ > 0.2; for larger values of 𝑛_𝑐_, the activated cell density in the initial sample is insufficient to yield dormancy break.

### Combined Control by Stochastic Activation and Quorum Rule

Since the stochastic or quorum rule-based mechanisms are by themselves unable to recreate the experimentally observed dynamics of FT1 expression and cell wall PD states, we next implemented a combination of both within the model. Implementing both mechanisms revealed a tradeoff landscape, which dictates how these mechanisms can influence plant’s behavior. Wildtype data for the cellular FT1 expression and the number of cell walls with open PDs are shown in **Figure 4A and C** for comparison to the simulated tissue landscape. First, looking at the effects of local coordination, the probability of stochastic activation is held constant (𝑝_𝑐_ = 0.0009), and the 𝑛_𝑐_ threshold is varied (**Figure 4B** for simulated gene expression and **Figure 4D** for simulated cell wall PD openness). These results corroborate those of **Figure 3**, where in both cases show phase transition behavior at 𝑛_𝑐_ > 0.2. When both mechanisms are present this transition signals the overtaking of the stochastic activation, making its contribution negligible in a continuous cold condition. Holding time constant (672hrs), 𝑝_𝑐_ and 𝑛_𝑐_ are varied as seen in **Figure 4E and F**, we find when 𝑛_𝑐_ < 0.2 the quorum rule becomes dominant, rendering the effects of the stochastic activation negligible. Conversely, for 𝑛_𝑐_ > 0.4, the stochastic activation becomes dominant over the quorum rule. The transition zone where both quorum rule and a stochastic mechanism have significant contributions occurs in the range 0.2 ≲ 𝑛_𝑐_ ≲ 0.4. Simulated cellular FT1 expression is a direct result of these two parameters, while simulated cell wall permeability dynamics (i.e., PD openness) are an effect of the cellular FT1 expression. These general trends are evident with the cell wall PDs opening rates trending behind the FT1 expression rates. The cell wall PD opening rates trending behind the FT1 expression rates is consistent with FT1 expression being necessary for PD opening as shown in experiments [41]. Importantly, these observations suggest conditions (a region of parameter space) under which the stochastic activation mechanism is dominant over quorum rule and vice-versa, and where both mechanisms may act in combination.

**Figure 4.**
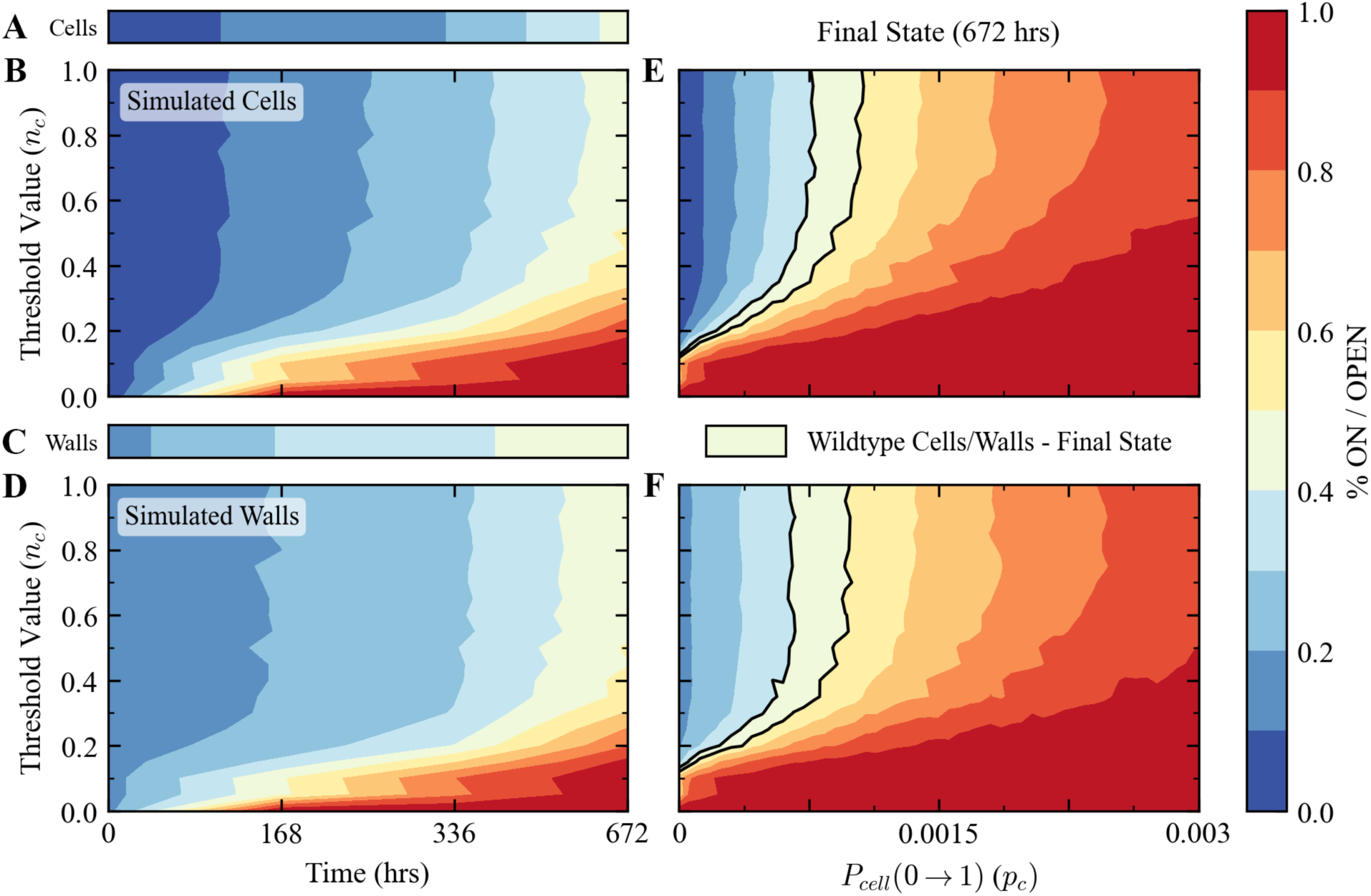
Results of the dynamic model highlighting the trade-off of stochastic versus collective coordination. Potential mechanisms for the activation of FT1 in continuous cold include stochastic activation and a quorum rule. 𝑛_𝑐_ is the threshold for quorum rule decision-making, corresponding to the percentage of neighbors required to be expressing FT1 for a cell to also begin expressing. The tissue template utilized is the 2-dimensional SAM template of the hybrid aspen meristematic tissue. **Top row:** Simulated gene expression levels (nodes). **Bottom row:** percentage of PDs open (edges). **Left column:** 𝑝_𝑐_ = 0.0009 and variable 𝑛_𝑐_ over simulated continuous cold treatment. Corresponding experimental data for expression levels are shown above the plots. **Right column:** Gene expression and cell wall PD openness at four weeks continuous cold treatment; FT1 expression levels and cell wall PD openness at four weeks continuous cold is outlined in black.

Comparing experimental data to the simulated data, we find the model is under-constrained i.e., there exist numerous combinations of parameters that could potentially reproduce the limited experimental data. Furthermore, we cannot entirely rule out the possibility that selecting alternative mechanisms, such as diffusion, could potentially yield better results. However, it is unlikely that other mechanistic assumptions would yield tighter constraints on a model, as our rules are already tightly coupled to known biological mechanism. One interesting avenue of future research could include generating high resolution FT1 expression data, and PD analysis data to further refine and constrain model parameters (a very costly process due to time and labor and sample limitations for slow growing aspen). With current data, to constrain 𝑝_𝑐_ and 𝑛_𝑐_ further, and to explore how multi-level constraints might lead to more informative models, we next consider population-level data capturing the statistics of bud-breaking across populations of hybrid aspen trees.

### Multiscale Modeling Including Population Behavior Can Constrain Tissue-Level Mechanisms

Plant populations across species commonly show bet-hedging behaviors [1], which include variable response to the same stimuli [23]. Among these, hybrid aspen trees also display bet hedging like behavior during dormancy release [43]. In populations of genetically identical hybrid aspen trees, bet hedging like response is observed in the heterogenous timing of bud break when constant cold is used to break dormancy. Moreover, this heterogeneity in bud break timing is further enhanced when variable cold is used for dormancy release [43]. Since bud break requires dormancy release, experimental data for bud break timing in a population of genetically identical hybrid aspen trees was compared to simulated populations defined by different combinations of 𝑝_𝑐_and 𝑛_𝑐_to identify which combination(s) best recreate both the population-level and cellular-level dynamics of FT1 expression. **Figure 4** displays results for the cellular level FT1 expression rates and cell wall PD openness rates, used to inform simulations of populations. Simulated populations were created utilizing the same tissue template and same random initial tissue state (1% of ON cells, 16.5% of cell walls have OPEN PDs). These simulated individuals represent the genetically identical plants used in the experiment. The simulated time interval is 960hrs (40 days) to match experimental protocols for population studies. **Figure 5** shows the population-level behaviors of the population (𝑁 = 25) and five simulated populations (𝑁 = 100) defined using different FT1 activation dynamics and parameter 𝑝_𝑐_ and 𝑛_𝑐_ values.

**Figure 5.**
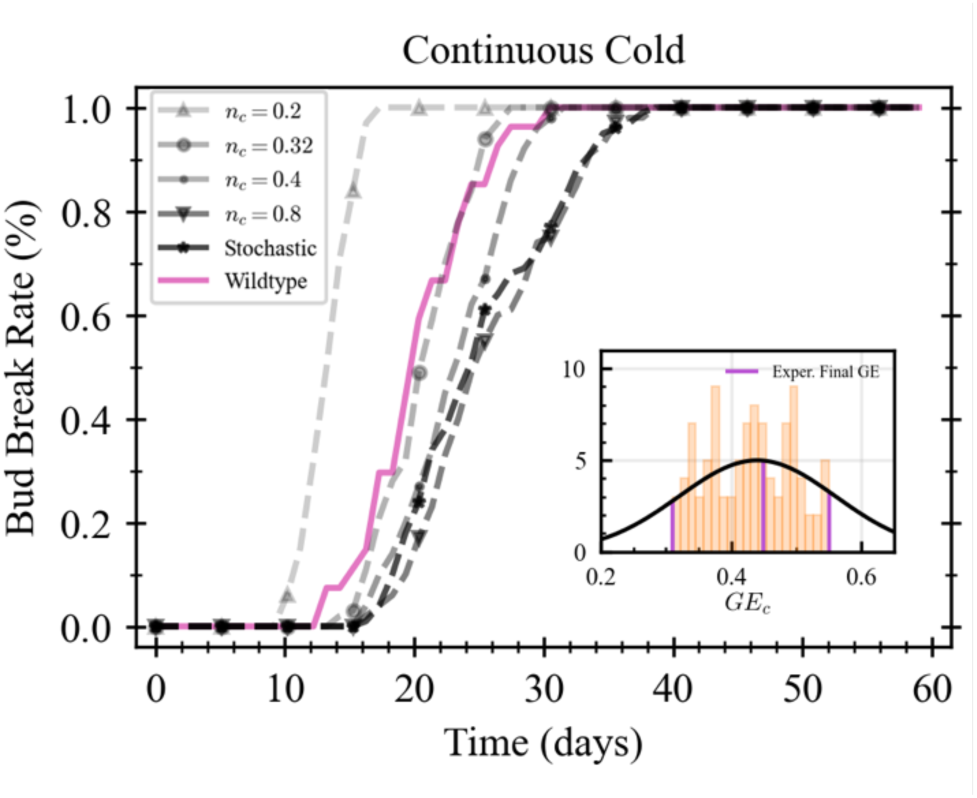
Population-level behaviors constrain 𝒏_𝒄_ in continuous cold treatment. “Simulated” denotes a wildtype tissue network model with 1% of nodes on and 16.5% of edges open at 𝑡 = 0. “Stochastic” is a model where only random activation of FT1 in response to cold exists, and no cellular coordination exists; the other four have both dynamics with differing 𝑛_𝑐_. The population with both mechanisms (stochastic activation 𝑝_𝑐_ = 0.0009 and quorum rule 𝑛_𝑐_ = 0.32) present showed the least error of all the simulated parameters in continuous cold (𝑀𝐴𝐸 = 0.016), see also **Supplement 6**.

In the experimental conditions, dormancy break is determined by visual inspection of the terminal bud for the necessary signs of growth. For simulated individuals, visual inspection is not possible; therefore, a critical threshold of total FT1 gene expression (𝐺𝐸_𝑐_,) is necessary to simulate bud dormancy breakage since FT1 expression is mechanistically responsible for dormancy breakage and PD opening. Under experimental conditions, dormancy release showed a correlation to FT1 expression between 0.31 ≤ 𝐺𝐸 ≤ 0.55 from 3 biological replicates (𝑁 = 3) exposed to continuous cold conditions. For each simulated individual plant tissue, 𝐺𝐸_𝑐_ was randomly selected from a truncated Gaussian created from the 3 replicates (𝜇 = 0.44, 𝜎 = 0.12), **Figure 5**. A Gaussian distribution allows us to work with the limited data available and is consistent with population level variation across diverse living systems often take a Gaussian form [19]. Additional distributions (single-value and uniform) for 𝐺𝐸_𝑐_ are considered in **Supplement 6**. We truncate the distribution to more closely match experimental gene expression data combined with visual inspection of wildtype individuals, which indicates some minimal amount of FT1 expression must be present for dormancy to break. Conversely, once enough FT1 is present, regardless of inter-individual variation in gene expression threshold, all individuals would break dormancy. In other words, while some individuals could break dormancy with a relatively lower expression of FT1, there exists a point where all individuals will break dormancy, thus informing an upper bound.

When stochastic activation of FT1 is the sole mechanism in the simulated population, the population-level behavior in the continuous cold reflects bet-hedging trends seen in the wildtype experiments, though slightly shifted to the right. When a quorum rule coordination is included the overall trend approaches that of the experimental data as 𝑛_𝑐_ decreases, with a best fit at 𝑛_𝑐_∼0.32. This suggests that a quorum rule mechanism within cellular collectives could be providing additional regulation to improve accuracy in decision making necessary for population level behavior. Utilizing population-level behaviors shows how the parameter space presented in **Figure 4** can be constrained to a plausible set of parameters capable of reproducing the data at each scale of data (gene expression, cell wall PD openness, and population level bet hedging).

### Variable Temperatures and Termination of Bud Break

Constant cold points to the stochastic activation of FT1 being largely able to explain bud break heterogeneity. However, in nature, temperature fluctuates with warm spikes interspersed with cold. Hence, we reasoned that it would be essential to obtain a model that could better explain bud break heterogeneity under such fluctuating conditions. Experimental data suggests that in contrast with constant cold, under fluctuating cold, warm spikes can significantly attenuate FT1 induction [43]. These results suggest that any mechanism to fully capture dormancy regulation must also take the potential input of FT1 suppression (not just activation). Living systems are well known to have such shutdown mechanisms that provide error correction when conditions become undesirable [26; 28]. Therefore, we include this downregulation of FT1 in our model and chose a stochastic variable to represent it. In our model, at any given hour that the cells are exposed to warm temperatures, a cell will turn OFF with 𝑃_1_(0) = 𝑝_w_. Given 𝑝_𝑐_ = 0.0009 and 𝑛_𝑐_ = 0.32, wildtype individuals exposed to fluctuating temperatures (20hrs 4–8°C and 4hrs 20°C) [42], were utilized to select an appropriate 𝑝_w_. **Figure 6A**, shows the simulated individuals versus and the FT1 expression data (𝑝_w_ = 0.024, 𝑀𝐴𝐸 = 0.0008), supporting that the shutdown dynamic of FT1 could be dictated by a stochastic process. Expanding to consider the population behavior of a fluctuating temperature regime (22hrs cold, 2hrs warm per day), **Figure 6B**, dormancy does break but at later timepoints than the individuals in constant cold (see **Figure 5)**. First, what this shows is that when only stochastic FT1 activation is considered, in contrast with constant cold temperatures, no individual breaks dormancy in variable cold conditions. The addition of cellular coordination adds necessary activation of FT1 expression in variable temperatures to ensure dormancy can break even under these conditions. Given the best fit from the continuous cold condition (𝑝_𝑐_ = 0.0009, 𝑛_𝑐_ = 0.32, 𝑝_w_ = 0.024) the simulated population shows a mean average error between simulated and experimental individuals of 0.05. To simulate the resultant bet hedging behavior in variable conditions all dynamics so far outlined are necessary.

**Figure 6.**
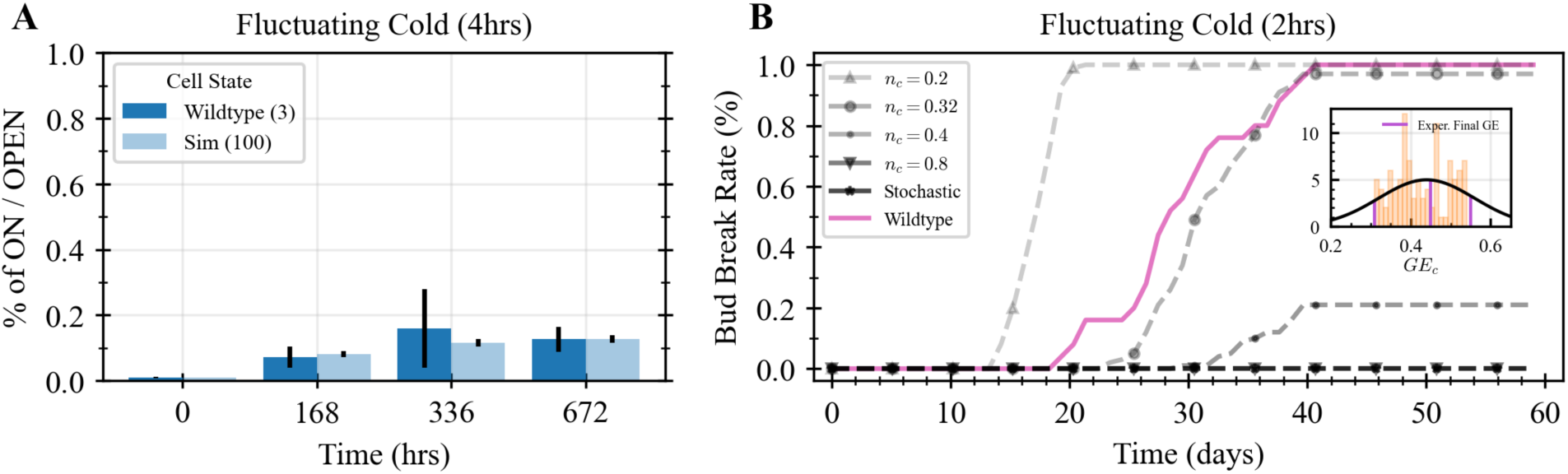
Variable Temperature Conditions effects on Individuals and Populations. **A)** Individual gene expression rates for wildtype and simulated individuals in variable temperatures 20hrs cold temperatures 4hrs warm temperatures. Initial state (𝑡 = 0) of wildtype samples has 1% of cells expressing FT1 and 16.5% of cell walls open; therefore, the simulated tissues began with 1% of nodes ON and 16.5% of edges OPEN, matching experimental initial conditions. If downregulation of FT1 is modelled via a stochastic variable the probability of any given cell turning OFF in the presence of warm temperatures was found to be 𝑝_*w*_ = 0.024 (𝑀𝐴𝐸 = 0.0008), assuming in cold temperatures stochastic activation 𝑝_𝑐_ = 0.0009 and a quorum rule 𝑛_𝑐_ = 0.32. **B)** 2hrs fluctuating cold treatment. Wildtype results are compared to five variations of simulated plants. Stochastic activation for FT1 is insufficient in varying temperatures conditions to result in any dormancy break. Some cellular coordination is necessary when environment variability is present. The simulated population considering all dynamics (stochastic activation, stochastic deactivation, and cellular coordination) reveal a reasonable fit to the wildtype population (𝑁 = 25) data with 𝑀𝐴𝐸 = 0.05.

The mechanisms that drive bet-hedging are not well understood. While many factors such as genotype and environmental differences would suffice in the wild, for the experimental data, individuals are genetically identical and are exposed to identical conditions, reducing the likelihood that the cause of bet-hedging like response is genotypic or environmental variation. Whatever the underlying mechanism, our model suggests that utilizing population-level behaviors can inform models of cellular-level behaviors, which in turn inform population level responses. It is therefore tempting to speculate that given the advantage of having heterogenous response in a population, cellular behaviors may have been selected for with higher-levels of organization constraining lower-levels, such that tissue decision making includes both bottom up and top down constraints.

## Discussion

In a multicellular organism, intercellular coordination creates the foundation for the complex behaviors that are key to growth and development. In contrast with animals, plants are advantageous for multiscale studies because they are sessile, their decision-making occurs on longer timescales, and their physical structure (at tissue/organ level) is readily mapped. Here we used the developmental transition from dormant state to release of dormancy in the terminal buds of hybrid aspen. The terminal bud dormancy release offers several advantages that are common to all higher plants, but additionally, during the switch to dormancy release there is no cell division or elongation [29]. This makes mapping the tissue and corresponding cellular interaction networks far easier than in other systems which have been utilized to study collective decision making. Finally, because the complexity of certain degrees of freedom (e.g., mobility) can be mitigated, or do not exist at all, greater isolation can occur to identify the components affecting decision-making, such as the exchange of information and connectivity between multiple levels of organization.

Here we used Boolean models to investigate collective decision making in buds to release dormancy in response to cold. Boolean models are frequently used to approximate the dynamics in complex systems when constructed using multiple organizational levels of information. Additionally, opening inquiry to consider intra-system informational feedback and utilizing additional levels can fill gaps in experimental data in the case where the necessary information at one level cannot be completely obtained. This is particularly advantageous in the case of buds where cellular resolution gene expression data cannot be easily obtained, and sampling (e.g., PD analysis) is often destructive. We show that the stochastic processes acting at the cellular level in a bud tissue can explain dormancy break for individual buds when subjected to constant cold – a somewhat idealized condition used in lab for uncovering genetic networks. However, stochastic processes cannot explain dormancy release when presented with conditions such as variable cold that buds typically experience in nature. This reveals a need for an additional regulatory component for robust regulation of bud dormancy release to explain individual as well as population-wide behaviors. For any population to display bet hedging, variation among individuals must exist. The plants studied herein are genetically identical individuals, in identical environments, and at the same developmental stage. The simulations replicating these experiments utilize these conditions and go so far as to make the simulated individuals of equal tissue size. Thus, both experimentally and theoretically much of the *intrinsic* variation is removed, which can typically contribute to variation. Here, we see the population-level response crucial for bet-hedging, an advantageous trait for fitness has presumably contributed to mechanisms (stochastic activation combined with quorum rule) at the cellular level. Furthermore, without a coordination dynamic within tissues, the robustness to error (i.e., being able to downregulate expression and return to dormancy) is reduced across all scales.

The model presented herein reveals wide reaching dynamical requirements to explain reported empirical data. However, there could exist other unidentified information and/or mechanisms that may be operating and contributing to the dormancy release process. For example, the model assumes instantaneous FT1 expression and immediate PD opening, omitting real-time processes like FT1 transcription, translation, and activation. Including these slower steps could alter parameter values and potentially lower the required 𝑛_𝑐_. The model also assumes a quorum-rule coordination among cells, though this remains to be experimentally demonstrated. Nevertheless, FT1 protein has been shown to move via PD in *Arabidopsis* as well FT1 induction of phytohormone gibberellic acid (GA) which has also been shown to move via PD [32]. This, combined with demonstration of both FT1 and GA in PD opening in buds, makes this mechanism plausible. Additionally, accounting for diffusion mechanisms and dynamic FT1 regulation, such as activation in cold conditions, suppression in warm, and PD closure, also enhances model precision and provides a more complete understanding of plant decision-making in response to environmental cues. Importantly, incorporating these potential mechanisms not only improves model predictions but crucially provides testable hypotheses for future experiments. Whereas the genetic networks underlying dormancy release are well described, we use this knowledge to reveal plausible collective decision making involved across multiple scales that was not known before.

## Acknowledgements

We would like to thank the hybrid aspen trees, without which this work would not be possible. Furthermore, we would like to thank the expertise and technical support of Dr. Fabrice Cordelières and Dr. Guillaume Maucort for the acquisition the plasmodesmata data. We thank Dr. Theodore Pavlic, Dr. Douglas G. Moore and Dr. Louie Slocombe for input on model development and improvement, and Eyal Segal for writing inspiration. This work was funded by Human Frontiers Science Program under Award No. RGP002/2020.

## Materials and Methods

For our model system the data acquired includes cellular tissue structure (including PD state distributions at cell interfaces), gene expression rates for FT1 protein in cellular tissue, dormancy states of individuals under study, and time of bud break for all individuals within a population [43]. Data concerning dormancy state and bud break are acquired by visual inspection of the plant and therefore are gathered without effect on the individual sample [43]. However, gene expression, PD state determination, and tissue organization can each only be acquired through measurement techniques that destroy the sample under examination; this excludes the possibility of using the same sample to collect these data. Gene expression, via RNA isolation and PCR sequencing, provides the total amount of FT1 present in each sample, but with no specificity as to which cells are expressing FT1. Furthermore, the states and distribution of cellular PDs are collected, via visual inspection, from different samples of the same tissue, but these samples are not the same ones from which the gene expression data was collected. Finally, tissue maps generated for simulation purposes are again different samples, because the techniques are different and each experiment is destructive to the sample. Therefore, the experimental datasets are uncoupled, elucidating a need for a biologically informed link, and prior expert knowledge of potential mechanisms, to inform model construction on the effects of the gene state on the cell wall state and vice versa. Recent work has shown the connection between the expression of FT1 and the opening of PD, i.e., for a PD to open, a cell must be expressing FT1 [42]. FT1’s direct link to PD state therefore allows the different tissue samples (provided from the same individual) to be coupled.

### Experimental Setup

The experimental setup focused on the transition of plants from dormancy to bud break in four different temperature regimes and encompassed both individual plants and plant populations. The temperature regimes were selected based on the conditions that *Populus spp.* requires for dormancy break to occur. Dormancy is initialized by simulating 10 – 11 weeks of short days. Continuous cold treatment is where, for 672hrs (28dys), the plants were placed in 4°C. This is sufficient to result in dormancy being broken and active growth occurring. Two fluctuating cold treatments where the plants were exposed to the same total number of hours at 4°C as in continuous cold treatment (672) but with additional intervals of warm temperatures of 20°C added, 2hrs or 4hrs daily, resulting in durations of 30 or 33 days respectively. 4hrs of warm temperatures is sufficient for dormancy not to be broken. Studies on small sets of individuals (3 ≤ 𝑁 ≤ 7) provides the gene expression and PD activity data for continuous cold and 4hrs fluctuating cold. While a different set of individuals (12 ≤ 𝑁 ≤ 27) provides population-level bud break behaviors for continuous cold, 2 hr, and 4 hr fluctuating cold treatments. In addition, detailed tissue mapping of the cells within the meristematic tissue was performed to make representative tissue networks. For a complete experimental set-up, see [43].

Pandey et al., 2025 reported that gene expression data was acquired for the expression of the Flowering Locus 1 (FT1) gene via RNA isolation and PCR sequencing. FT1 expression ratio was treated equal to the likely ratio found across the entire individual. Rates were sampled in the dormant state, at 168h, 336h, and 672h for continuous cold and 4hrs fluctuating cold treatments. PD activity was acquired from one individual in the continuous cold treatment and one in the 4hrs fluctuating cold. Six cross-sectional samples were acquired from each individual in the dormant state, at 168h, 336h, 504h, and 672h. These samples were then treated and imaged using transmission electron microscopy. From there, individual PD states were identified by visual inspection as either opened or closed, with cellular networks being constructed with cell walls characterized by the percentage of PDs that were opened. Finally, population-level behavior focused only on determining the days in which plants within the study set broke bud dormancy; this was done by visual inspection of the terminal buds. Population sizes for the tested temperate are 𝑁 = 27 for continuous cold, 𝑁 = 25 for 2hrs fluctuating cold, and 𝑁 = 12 for 4hrs fluctuating cold.

### Boolean Model for Cellular Dynamics

*Populus spp.* is primarily influenced by external temperature for the breaking of bud dormancy. Therefore, the dynamics are modeled in response to external temperature. Starting with response to cold temperatures, the hypothesis is that a cellular coordination mechanism is necessary for accurate activation of cells. Therefore, the state update rule for a given node, 𝑠, and conditioned on the set of its nearest neighbor node states, 𝑠̅, and their corresponding edge states, 𝜌̅.

The state update rule for a given cell, 𝑠, is:

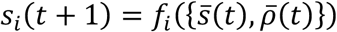

Where the function comprises a random probability of switching to the ON state and a cellular coordination mechanism in the form of a quorum rule:

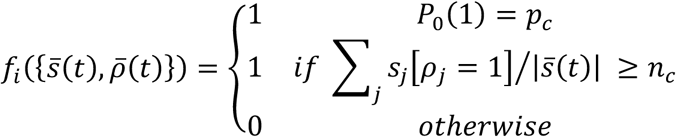

This translates to: if the cell does not randomly begin expressing FT1 (with a probability 𝑝_𝑐_) in the presence of cold temperatures, then if the ratio of its neighboring cells that are expressing and have an OPEN cell wall to the total number of neighbors is greater than 𝑛_𝑐_, the cell will begin to express FT1 at the next timestep; if not, it will remain in an off state. Neither 𝑝_𝑐_ or 𝑛_𝑐_ are known *a priori,* they will be identified based on validation with experimental data.

Due to the now-known connection between the opening of PDs and the expression of FT1, the state update rule for a cell wall can be conditioned on the cells in which it connects. Once again, this is in response to cold temperatures.

For a given cell wall, 𝜌, the edge update rule is:

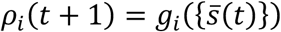

Where the function capturing the cell wall (edge) update is:

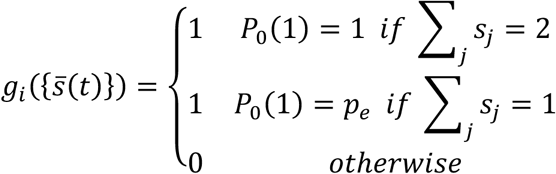

That is, if a cell wall borders two cells expressing FT1, the cell wall opens else if only one cell is expressing FT1, there is a random chance, 𝑝_𝑒_, for the PD to open, and if neither cell expresses FT1, the PD does not open.

In response to warm temperatures, the update function for cells that are becomes:

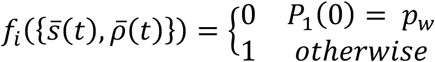

Due to limitations in both the understanding of the study system and availability of data, the following assumptions are within the model:

- Cells ON will not turn OFF in cold temperatures
- Cell OFF will not turn ON in warm temperatures
- Cell walls once OPENED cannot be CLOSED

## Author Contributions

**Hannah Dromiack**: model conceptualization; methodology (lead); formal analysis (lead); and writing – original draft preparation (lead). **Swanand Khanapurkar**: methodology (supporting), conceptualization (supporting). **Rachel Phillips**: methodology (supporting). **Tatiana de Souza Moraes**: investigation (equal); writing – review and editing (supporting). **Gwendolyn Davis**: investigation (equal); writing – review and editing (supporting). **Shashank Pandey**: investigation (equal). **Bibek Aryal**: investigation (equal). **Aswin Nair**: investigation (equal). **George Bassel**: project conceptualization; funding acquisition (equal); supervision (supporting); writing – review and editing (supporting). **Emmanuelle Bayer**: project conceptualization; funding acquisition (equal); supervision (supporting); writing – review and editing (supporting). **Rishikesh Bhalerao**: project conceptualization; funding acquisition (equal); supervision (supporting); writing – review and editing (supporting). **Sara Walker**: project conceptualization; funding acquisition (equal); supervision (lead); writing – review and editing (lead).

## References

[1] K. Abley, R. Goswami, J.C.W. Locke, Bet-hedging and variability in plant development: seed germination and beyond, Philosophical Transactions of the Royal Society B: Biological Sciences. 379 (2024). 10.1098/rstb.2023.0048.

[2] P.W. Anderson, More Is Different: Broken symmetry and the nature of the hierarchical structure of science., Science. 177 (1972) 393–396. 10.1126/science.177.4047.393.

[3] D. André, A. Marcon, K.C. Lee, D. Goretti, B. Zhang, N. Delhomme, M. Schmid, O. Nilsson, FLOWERING LOCUS T paralogs control the annual growth cycle in *Populus* trees, Current Biology. 32 (2022) 2988–2996.e4. 10.1016/j.cub.2022.05.023.

[4] A. Angel, J. Song, C. Dean, M. Howard, A Polycomb-based switch underlying quantitative epigenetic memory, Nature. 476 (2011) 105–108. 10.1038/nature10241.

[5] Y.M. Bar-On, R. Phillips, R. Milo, The biomass distribution on Earth, Proceedings of the National Academy of Sciences. 115 (2018) 6506–6511. 10.1073/pnas.1711842115.

[6] G.W. Bassel, To Grow or not to Grow?, Trends in Plant Science. 21 (2016) 498–505. 10.1016/j.tplants.2016.02.001.

[7] A. Berdahl, C.J. Torney, C.C. Ioannou, J.J. Faria, I.D. Couzin, Emergent Sensing of Complex Environments by Mobile Animal Groups, Science. 339 (2013) 574–576. 10.1126/science.1225883.

[8] I.D. Couzin, J. Krause, R. James, G.D. Ruxton, N.R. Franks, Collective Memory and Spatial Sorting in Animal Groups, Journal of Theoretical Biology. 218 (2002) 1–11. 10.1006/jtbi.2002.3065.

[9] F. Cucker, S. Smale, Emergent Behavior in Flocks, IEEE Transactions on Automatic Control. 52 (2007) 852–862. 10.1109/tac.2007.895842.

[10] G.V. Davis, T. de Souza Moraes, S. Khanapurkar, H. Dromiack, Z. Ahmad, E.M. Bayer, R.P. Bhalerao, S.I. Walker, G.W. Bassel, Toward uncovering an operating system in plant organs, Trends in Plant Science. 29 (2024) 742–753. 10.1016/j.tplants.2023.11.006.

[11] C. Detrain, J.-L. Deneubourg, J.M. Pasteels, Decision-making in foraging by social insects, in: Information Processing in Social Insects, Birkhäuser Basel, 1999: pp. 331–354. 10.1007/978-3-0348-8739-7_18.

[12] S. Duran-Nebreda, G.W. Bassel, Plant behaviour in response to the environment: information processing in the solid state, Philosophical Transactions of the Royal Society B: Biological Sciences. 374 (2019) 20180370. 10.1098/rstb.2018.0370.v

[13] G.F.R. Ellis, Top-down causation and emergence: some comments on mechanisms, Interface Focus. 2 (2011) 126–140. 10.1098/rsfs.2011.0062.

[14] P. Erdős, A. Rényi, On random graphs. I., Publicationes Mathematicae Debrecen. 6 (2022) 290–297. 10.5486/pmd.1959.6.3-4.12.

[15] J.K. Feibleman, Theory of Integrative Levels, The British Journal for the Philosophy of Science. 5 (1954) 59–66. 10.1093/bjps/v.17.59.

[16] G.P. Figueredo, T.V. Joshi, J.M. Osborne, H.M. Byrne, M.R. Owen, On-lattice agent-based simulation of populations of cells within the open-source Chaste framework, Interface Focus. 3 (2013) 20120081. 10.1098/rsfs.2012.0081.

[17] J. Flack, Life’s Information Hierarchy, in: From Matter to Life, Cambridge University Press, 2017: pp. 283–302. 10.1017/9781316584200.012.

[18] S. Fox, P. Southam, F. Pantin, R. Kennaway, S. Robinson, G. Castorina, Y.E. Sánchez-Corrales, R. Sablowski, J. Chan, V. Grieneisen, A.F.M. Marée, J.A. Bangham, E. Coen, Spatiotemporal coordination of cell division and growth during organ morphogenesis, PLOS Biology. 16 (2018) e2005952. 10.1371/journal.pbio.2005952.

[19] S.A. Frank, The common patterns of nature, Journal of Evolutionary Biology. 22 (2009) 1563–1585. 10.1111/j.1420-9101.2009.01775.x.

[20] M. Gianella, K.J. Bradford, F. Guzzon, Ecological, (epi)genetic and physiological aspects of bet-hedging in angiosperms, Plant Reproduction. 34 (2021) 21–36. 10.1007/s00497-020-00402-z.

[21] D.M. Gordon, Ants at work: How an insect society is organized, Norton, New York [u.a.], 2000.

[22] D.M. Gordon, Measuring collective behavior: an ecological approach, Theory in Biosciences. 140 (2019) 353–360. 10.1007/s12064-019-00302-5.

[23] A.J. Grimbergen, J. Siebring, A. Solopova, O.P. Kuipers, Microbial bet-hedging: the power of being different, Current Opinion in Microbiology. 25 (2015) 67–72. 10.1016/j.mib.2015.04.008.

[24] A.A. Hagberg, D.A. Schult, P.J. Swart, Exploring Network Structure, Dynamics, and Function using NetworkX, in: Proceedings of the 7th Python in Science Conference, SciPy, 2008: pp. 11–15. 10.25080/tcwv9851.

[25] S. Jansson, C.J. Douglas, Populus: A Model System for Plant Biology, Annual Review of Plant Biology. 58 (2007) 435–458. 10.1146/annurev.arplant.58.032806.103956.

[26] R.W. Jones, Principles of biological regulation: An introduction to feedback systems, Academic Press, New York, 1973.

[27] E. Karahanna, J.R. Evaristo, M. Srite, Levels of Culture and Individual Behavior: An Investigative Perspective, Journal of Global Information Management. 13 (2005) 1–20. 10.4018/jgim.2005040101.

[28] H. Kitano, Biological robustness, Nature Reviews Genetics. 5 (2004) 826–837. 10.1038/nrg1471.

[29] T. Kean-Galeno, D. Lopez-Arredondo, L. Herrera-Estrella, The Shoot Apical Meristem: An Evolutionary Molding of Higher Plants, International Journal of Molecular Sciences. 25 (2024) 1519. 10.3390/ijms25031519.

[30] J. Kolasa, S.T.A. Pickett, Ecological systems and the concept of biological organization, Proceedings of the National Academy of Sciences. 86 (1989) 8837– 8841. 10.1073/pnas.86.22.8837.

[31] Z.P. Li, A. Paterlini, M. Glavier, E.M. Bayer, Intercellular trafficking via plasmodesmata: molecular layers of complexity, Cellular and Molecular Life Sciences. 78 (2020) 799–816. 10.1007/s00018-020-03622-8.

[32] L. Liu, C. Liu, X. Hou, W. Xi, L. Shen, Z. Tao, Y. Wang, H. Yu, FTIP1 Is an Essential Regulator Required for Florigen Transport, PLoS Biology. 10 (2012) e1001313. 10.1371/journal.pbio.1001313

[33] A. Lloret, M.L. Badenes, G. Ríos, Modulation of Dormancy and Growth Responses in Reproductive Buds of Temperate Trees, Frontiers in Plant Science. 9 (2018). 10.3389/fpls.2018.01368.

[34] E. Mallon, S., N. Franks, Individual and collective decision-making during nest site selection by the ant *Leptothorax albipennis*, Behavioral Ecology and Sociobiology. 50 (2001) 352–359. 10.1007/s002650100377.

[35] H.H. McAdams, A. Arkin, Stochastic mechanisms in gene expression, Proceedings of the National Academy of Sciences. 94 (1997) 814–819. 10.1073/pnas.94.3.814.

[36] P.A.M. Mediano, F.E. Rosas, A.I. Luppi, R.L. Carhart-Harris, D. Bor, A.K. Seth, A.B. Barrett, Towards an extended taxonomy of information dynamics via Integrated Information Decomposition, arXiv. (2021). 10.48550/ARXIV.2109.13186.

[37] S. Milgram, Group pressure and action against a person., The Journal of Abnormal and Social Psychology. 69 (1964) 137–143. 10.1037/h0047759.

[38] N. Miller, S. Garnier, A.T. Hartnett, I.D. Couzin, Both information and social cohesion determine collective decisions in animal groups, Proceedings of the National Academy of Sciences. 110 (2013) 5263–5268. 10.1073/pnas.1217513110.

[39] W.L. Ng, B.L. Bassler, Bacterial Quorum-Sensing Network Architectures, Annual Review of Genetics. 43 (2009) 197–222. 10.1146/annurev-genet-102108-134304

[40] K.J. Niklas, The evolutionary biology of plants, [Nachdr.], Univ. of Chicago Pr., Chicago [u.a.], 1999.

[41] A.B. Novikoff, The Concept of Integrative Levels and Biology, Science. 101 (1945) 209–215. 10.1126/science.101.2618.209.

[42] S.K. Pandey, J.P. Maurya, B. Aryal, K. Drynda, A. Nair, P. Miskolczi, R.K. Singh, X. Wang, Y. Ma, T. de Souza Moraes, E.M. Bayer, E. Farcot, G.W. Bassel, L.R. Band, R.P. Bhalerao, A regulatory module mediating temperature control of cell-cell communication facilitates tree bud dormancy release, The EMBO Journal. 43 (2024) 5793–5812. 10.1038/s44318-024-00256-5.

[43] S.K. Pandey, T.S. Moraes, Aryal Bibek, A. Nair, P. Miskolczi, G. Maucort, F.P. Cordelières, G.V. Davis, H. Dromiack, S. Khanapurkar, S.I. Walker, G.W. Bassel, E.M. Bayer, R.P. Bhalerao, Variable Temperature Processing by Plasmodesmatal Network Regulates robust Bud Dormancy release, Accepted: Nature Communications (2025).

[44] J. Paulsson, Models of stochastic gene expression, Physics of Life Reviews. 2 (2005) 157–175. 10.1016/j.plrev.2005.03.003.

[45] S.C. Pratt, D.J.T. Sumpter, E.B. Mallon, N.R. Franks, An agent-based model of collective nest choice by the ant *Temnothorax albipennis*, Animal Behaviour. 70 (2005) 1023–1036. 10.1016/j.anbehav.2005.01.022

[46] J.H. Priestly, The Meristematic Tissues of the Plant, Biological Reviews. 3 (1928) 1–20. 10.1111/j.1469-185x.1928.tb00881.x.

[47] D. Pumain, ed., Hierarchy in Natural and Social Sciences, Springer Netherlands, 2006. 10.1007/1-4020-4127-6.

[48] P.C. Rogers, B.D. Pinno, J. Šebesta, B.R. Albrectsen, G. Li, N. Ivanova, A. Kusbach, T. Kuuluvainen, S.M. Landhäusser, H. Liu, T. Myking, P. Pulkkinen, Z. Wen, D. Kulakowski, A global view of aspen: Conservation science for widespread keystone systems, Global Ecology and Conservation. 21 (2020) e00828. 10.1016/j.gecco.2019.e00828.

[49] M. Salahshour, I.D. Couzin, Allocentric Flocking, bioRxiv. (2025). 10.1101/2025.01.27.634610.

[50] G.C. Santos, Upward and Downward Causation from a Relational-Horizontal Ontological Perspective, Axiomathes. 25 (2014) 23–40. 10.1007/s10516-014-9251-x.

[51] D.J.T. Sumpter, S.C. Pratt, Quorum responses and consensus decision making, Philosophical Transactions of the Royal Society B: Biological Sciences. 364 (2008) 743–753. 10.1098/rstb.2008.0204.

[52] J.W. Valentine, C.L. May, Hierarchies in biology and paleontology, Paleobiology. 22 (1996) 23–33. 10.1017/s0094837300015992.

[53] T. Vicsek, A. Czirók, E. Ben-Jacob, I. Cohen, O. Shochet, Novel Type of Phase Transition in a System of Self-Driven Particles, Physical Review Letters. 75 (1995) 1226–1229. 10.1103/physrevlett.75.1226.

[54] P.K. Visscher, Group Decision Making in Nest-Site Selection Among Social Insects, Annual Review of Entomology.52 (2007) 255–275. 10.1146/annurev.ento.51.110104.151025.

[55] T. Watanabe, Y. Takahashi, Tissue morphogenesis coupled with cell shape changes, Current Opinion in Genetics and Development. 20 (2010) 443–447. 10.1016/j.gde.2010.05.004.

[56] J.L. Whited, M. Levin, Bioelectrical controls of morphogenesis: from ancient mechanisms of cell coordination to biomedical opportunities, Current Opinion in Genetics and Development. 57 (2019) 61–69. 10.1016/j.gde.2019.06.014.

[57] D.P. Wickland, Y. Hanzawa, The FLOWERING LOCUS T/TERMINAL FLOWER 1 Gene Family: Functional Evolution and Molecular Mechanisms, Molecular Plant. 8 (2015) 983– 997. 10.1016/j.molp.2015.01.007.

[58] R. Worrell, European aspen (*Populus tremula L.*): a review with particular reference to Scotland I. Distribution, ecology and genetic variation, Forestry. 68 (1995) 93–105. 10.1093/forestry/68.2.93.

[59] A. Zafeiris, T. Vicsek, Why We Live in Hierarchies?: A Quantitative Treatise, Springer International Publishing, 2018. 10.1007/978-3-319-70483-8.

