## Supplement for "A Multiscale Model of Collective Decision-Making in Hybrid Aspen Tree Tissues Describes Bud-Dormancy Break"

### Supplement 1. Nomenclature

| | |
|---|---|
| $C$ | clustering coefficient |
| $C_k(g)$ | degree centrality |
| $d(u, v)$ | distance between some node $u$ and node $v$ |
| $\langle d(u, v) \rangle$ | average shortest path for all pairs $(u, v)$ |
| g | network |
| $i_e$ | percentage of OPEN edges at $t = 0$ |
| $i_n$ | percentage of ON nodes at $t = 0$ |
| $k$ | node degree |
| $max\ d(u, v)$ | network diameter, maximum path for all pairs $(u, v)$ |
| $n$ | $\sum_j s_j[\rho_j = 1]/\lvert\bar{s}(t)\rvert$ |
| $n_c$ | threshold value, for the node to change states $0 \rightarrow 1$ then $n \geq n_c$ |
| N | population size |
| $p_e$ | probability of an edge opening in cold temperatures, given one neighboring node is ON |
| $p_c$ | probability of a node stochastically turning ON in cold temperatures |
| $p_w$ | probability of a node stochastically turning OFF in warm temperatures |
| $p(k)$ | degree distribution |
| $r$ | degree assortativity |
| $\rho_i$ | $i^{th}$ edge |
| $\bar{\rho}$ | set of edges associated to neighboring cells of the $i^{th}$ cell |
| $s_i$ | $i^{th}$ node |
| $\bar{s}$ | set of neighboring nodes to the $i^{th}$ cell |
| CLOSED | $i^{th}$ edge with $\rho_i = 0$ |
| OFF | $i^{th}$ node with $s_i = 0$ |
| ON | $i^{th}$ node with $s_i = 1$ |
| OPEN | $i^{th}$ edge with $\rho_i = 1$ |

### Supplement 2. Network/Rule Visualization

#### Network Representation

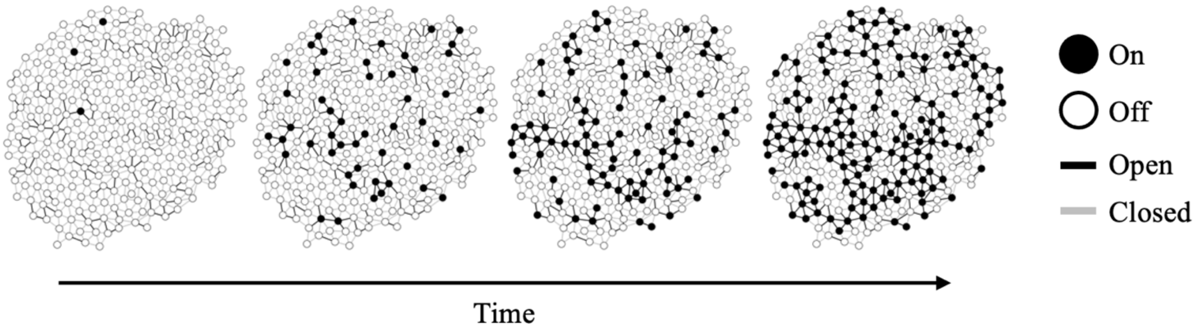


**Supplement Figure 1. Network representation over four weeks of 2D SAM tissue.** Cells are represented as nodes (white – not expressing FT1, black – expressing FT1) and cell walls as edges (grey – closed PDs, black – open PDs). Left to right: Dormant state 10 weeks short days, continuous cold at 1 week, 2 weeks, and 4 weeks.

#### Edge Update Rule

Edges can either be open or closed and that is represented as a binary variable [open = 1, close = 0]. At each timestep whether each edge $\rho$ opens or not is based on the state of the two cells it connects.

If both cells are ON

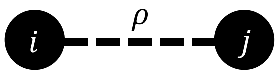


The cell wall will OPEN, this is because of the mechanism of PD opening is correlated to gene expression in the connected cells, therefore with both ON it is maximally effective.

If only one cell is ON

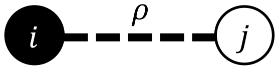


The cell wall will have a probability of OPENING equal to $\boldsymbol{p_e}$. This captures the idea that the cell wall can open because it is connected to a cell that is on.

If both cells are OFF

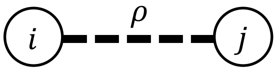


There is no mechanism for opening the PDs therefore the cell wall has no chance of opening.

#### Quorum Rule – Node Update Rule

At each timestep some node $s$ state is determined via the states of the neighboring cells and the state of the edges between those neighbors.

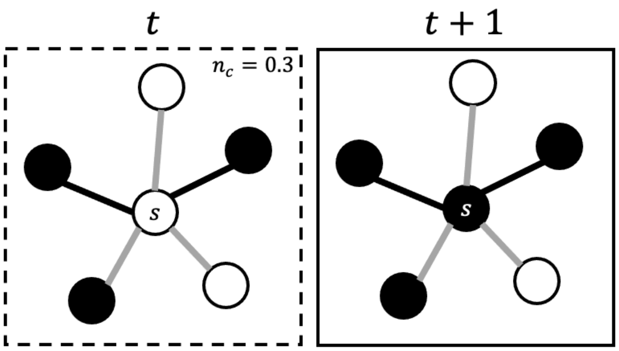


### Supplement 3. Temperature Treatments

Considering both a coordination dynamic and stochastic activation in the continuous cold treatment 100 simulated plants are compared to the experimental data, **Sup Figure 2**. $p_e$ was experimentally derived while $p_c$ and $p_w$ were determined by the model parameter sweeps.

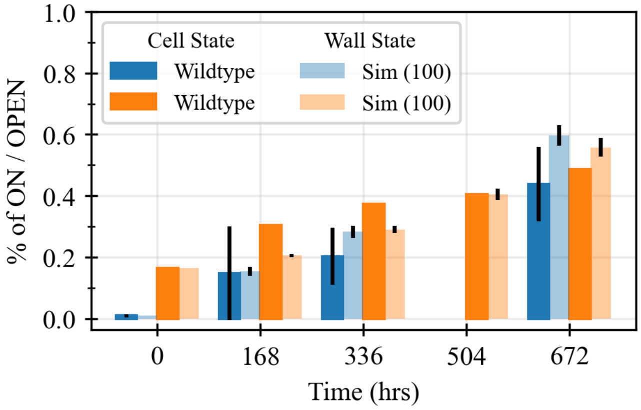


**Supplement Figure 2. Temperature Treatments.** Simulated continuous cold treatment. Initial conditions at $t = 0$ was determined via the dormant state, therefore 1% of nodes active and 16.5% of edges open.

### Supplement 4. Experimental Derived Quantities

As shown in the continuous warm treatment, portions of this model can be dictated via the experimentally derived quantities. For the stochastic (de)activation parameter $p_e$ is determined via exponential fits using least squares algorithm and $R^2$ values.

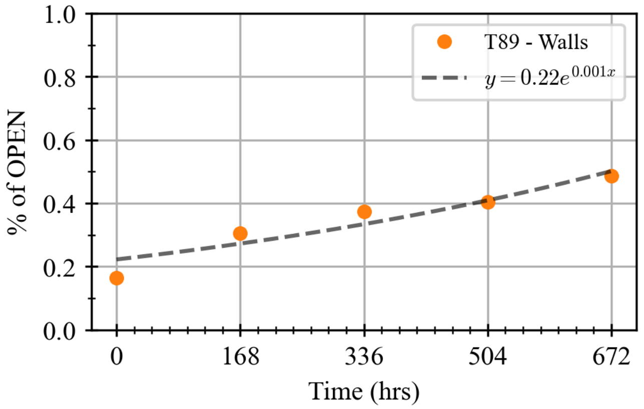


**Supplement Figure 3. Identifying $p_e$.** Rate of cell walls opening in cold temperatures as determined by an exponential fit ($R^2 = 0.89$).

Regardless of the temperature regime or dormancy there if observed activity within the tissue. For the dormant state, the initial number of edges OPEN, $i_e = 16.5\%$, is acquired from the 11wSD dormant state (Sample Set 1) comprised of 1 plant with 6 samples. The initial number of nodes ON, $i_n = 1\%$, is acquired from the 10 weeks short days dormant state comprised of 6 plants.

### Supplement 5. Expanded Network Analysis

Utilizing a standard toolbox of network statistics (average shortest path, clustering coefficient, network diameter, average degree centrality, degree assortativity, and degree distribution), we compared the plant tissue to connected Erdős Rényi (ER) graphs with a conserved distribution [Erdős and Rényi, 1959], **Sup Figure 4**. This standard suite of statistics was selected as they provide information concerning the overall structure of the tissue. The goal is to identify any meaningful features of the plant structure that may inform what possible decision-making strategies are utilized by these organisms in the process of dormancy breaking. An ensemble of 120 connected ER graphs was generated for each sample set; sample sets included six tissue samples; 20 ER graphs were generated for each network representation. Three primary conclusions are drawn from these, first that plant tissue is not a random but organized structure, this would agree with plants being organisms having evolved structures for specific functions [Niklas1999], **Sup Figure 4 top row and bottom row**. Second, many of the network statistics commonly used in scale-free networks are inadequate and uninformative for use in rigid topologies, **Sup Figure 4 middle row.** These inadequacies are due to this type of network topology not being driven by preferential attachment or connectiveness but intrinsic to the organism. A cell's neighbor is not one it chooses but is a result of other processes such as geometry, orientation of divisions and subsequent expansion. Thirdly, to determine the likely dynamics dictating the opening of PD, these results must be combined with externally gathered data, such as the mechanistic connection between the expression of FT1 and the opening of PDs.

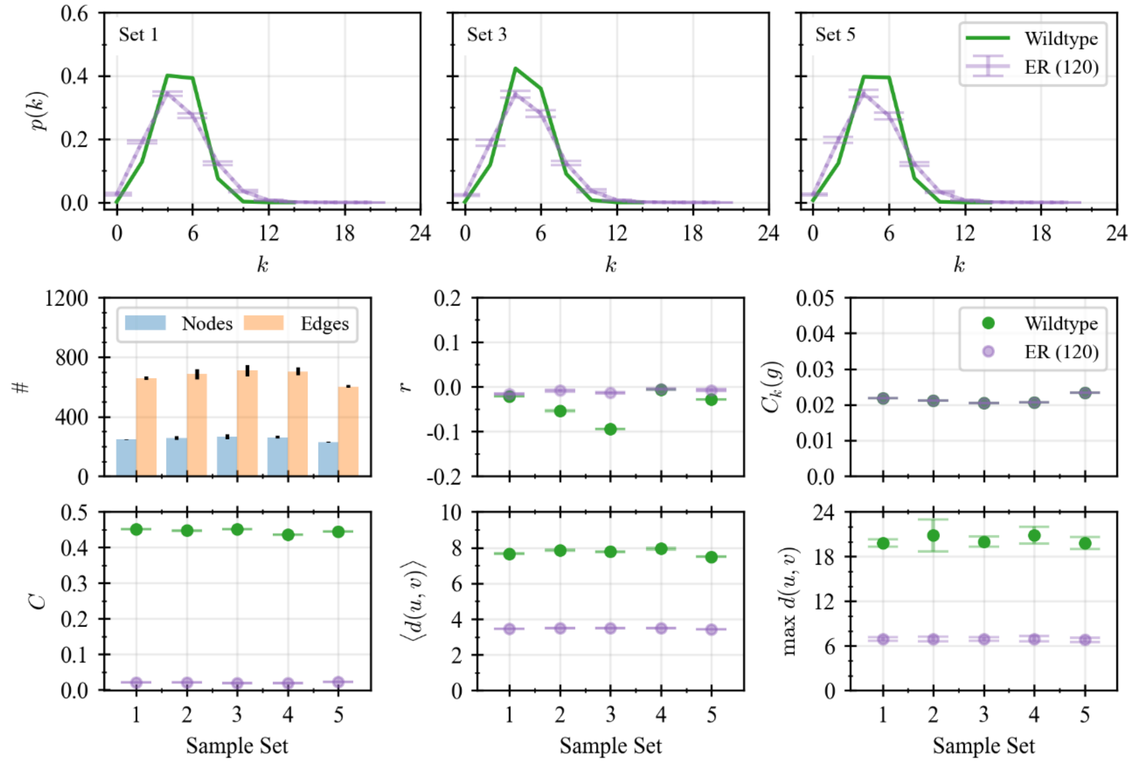


**Supplement Figure 4. Network Topology of PD Activity.** Top row: degree distribution, $p(k)$, of PD networks compared to connected Erdos Renyi (ER) networks in the dormant state, two weeks of continuous cold, and four weeks of continuous cold. While the general trend is similar, there is a flattening of the distribution in the random graphs that is significantly different from that of the wildtype networks. Middle row: Network characteristics (# of nodes and edges for each network), degree assortativity ($r$), degree centrality ($C_k(g)$). Due to the rigid topology of the plant tissue and connected ER networks, these measures provide no meaningful information. The network's connectivity is predetermined by evolution or randomization; thus, there is no preferential treatment based on connectivity. Bottom row: Clustering coefficient ($C$), average shortest path ($\langle d(u,v)\rangle$), and network diameter ($max\ d(u,v)$). The lattice-like structure of the aspen tissue is significantly different from that of a random topology, as seen by the bottom row. Longer path lengths exist in the wildtype because cells of the opposite end of the tissue cannot be direct neighbors to one another, as can happen in the ER network. Finally, more cluster triangles exist due to the lattice structure.

### Supplement 6. Population-level Behavior

**Table 1.** Mean Absolute Error values given a truncated gaussian $GE_c$ distribution for the simulations with $p_c = 0.0009$ and $p_w = 0.017$.

| | $n_c = 0.2$ | $n_c = 0.32$ | $n_c = 0.4$ | $n_c = 0.8$ | Stochastic |
|---|---|---|---|---|---|
| Continuous Cold | 0.117 | 0.016 | 0.048 | 0.093 | 0.082 |
| Fluctuating Cold | 0.215 | 0.029 | 0.219 | 0.487 | 0.485 |

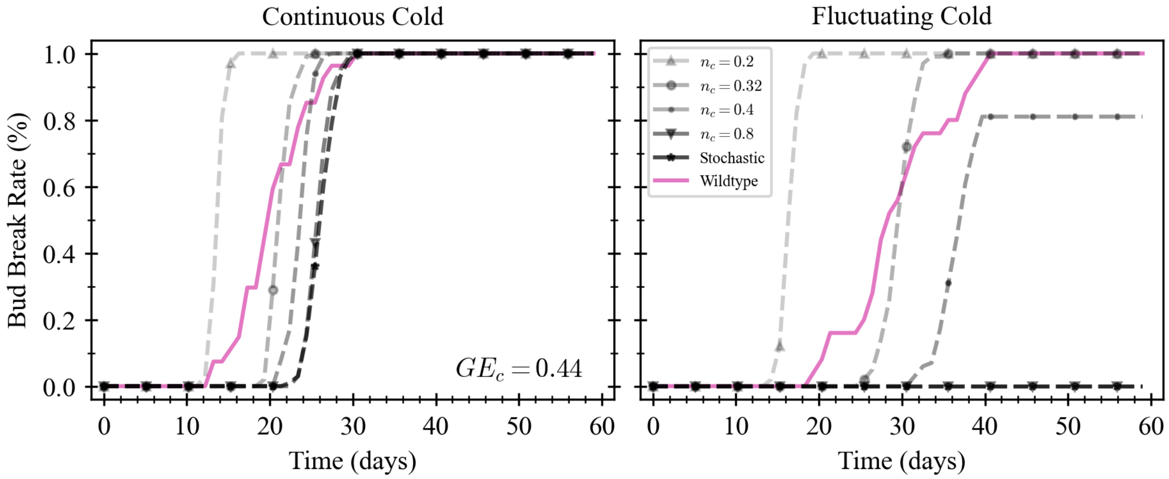


**Supplement Figure 5. Population Dynamics behaviors constrain $n_c$.** Simulated wildtype tissue with 16.5% of edges open at $t = 0$. Left: continuous cold treatment. Right: 2hrs fluctuating cold treatment. Wildtype results compared to five variations of simulated plants: Stochastic presents the model where only random activation of FT1 in response to cold exists, and no cellular coordination exists; the other four have both dynamics with differing $n_c$. A single value was selected for $GE_c$.

**Table 2.** Mean Absolute Error values given a *single value* $GE_c = 0.44$ for the simulations presented in Supplement Figure 5.

| | $n_c = 0.2$ | $n_c = 0.32$ | $n_c = 0.4$ | $n_c = 0.8$ | Stochastic |
|---|---|---|---|---|---|
| Continuous Cold | 0.109 | 0.044 | 0.065 | 0.094 | 0.100 |
| Fluctuating Cold | 0.215 | 0.056 | 0.189 | 0.504 | 0.504 |

$GE_c$ distribution was generated from the experimental gene expression data for individuals ($N = 3$) exposed to continuous cold conditions.

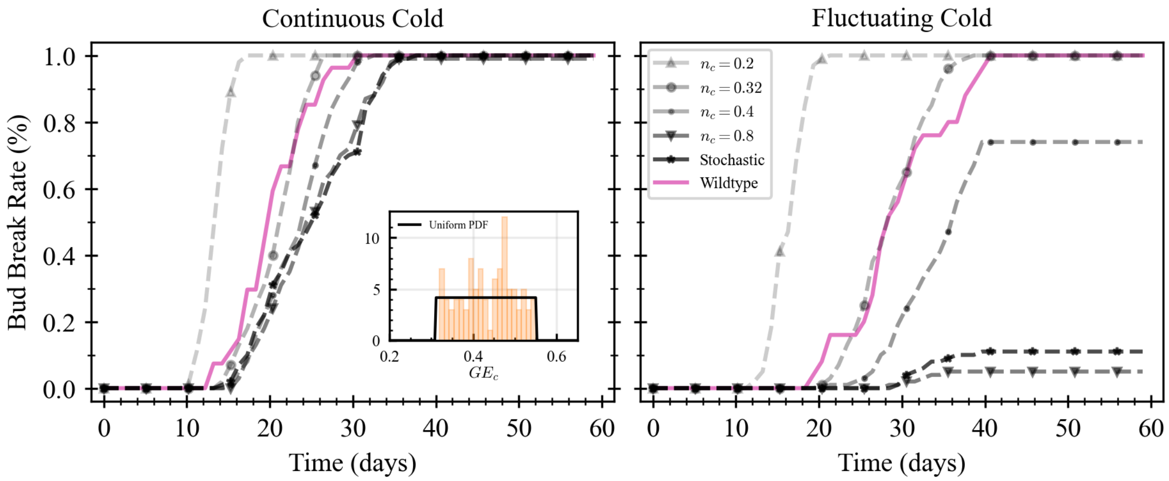


**Supplement Figure 6. Population Dynamics behaviors constrain $n_c$.** Simulated wildtype tissue with 16.5% of edges open at $t = 0$. Left: continuous cold treatment. Right: 2hrs fluctuating cold treatment. Wildtype results compared to five variations of simulated plants: Stochastic presents the model where only random activation of FT1 in response to cold exists, and no cellular coordination exists; the other four have both dynamics with differing $n_c$. A uniform distribution was selected for $GE_c$.

**Table 3.** Mean Absolute Error values given a uniform $GE_c$ distribution for the simulations with $p_c = 0.0009$ and $p_w = 0.017$.

| | $n_c = 0.2$ | $n_c = 0.32$ | $n_c = 0.4$ | $n_c = 0.8$ | Stochastic |
|---|---|---|---|---|---|
| Continuous Cold | 0.115 | 0.020 | 0.051 | 0.085 | 0.079 |
| Fluctuating Cold | 0.218 | 0.025 | 0.177 | 0.481 | 0.455 |
